## Supplementary Information for "Benchmarking and Optimization of Methods for the Detection of Identity-By-Descent in High-Recombining *Plasmodium falciparum* Genomes"

Guo *et al.*

### 1 Supplementary Figures

**S1 Fig**

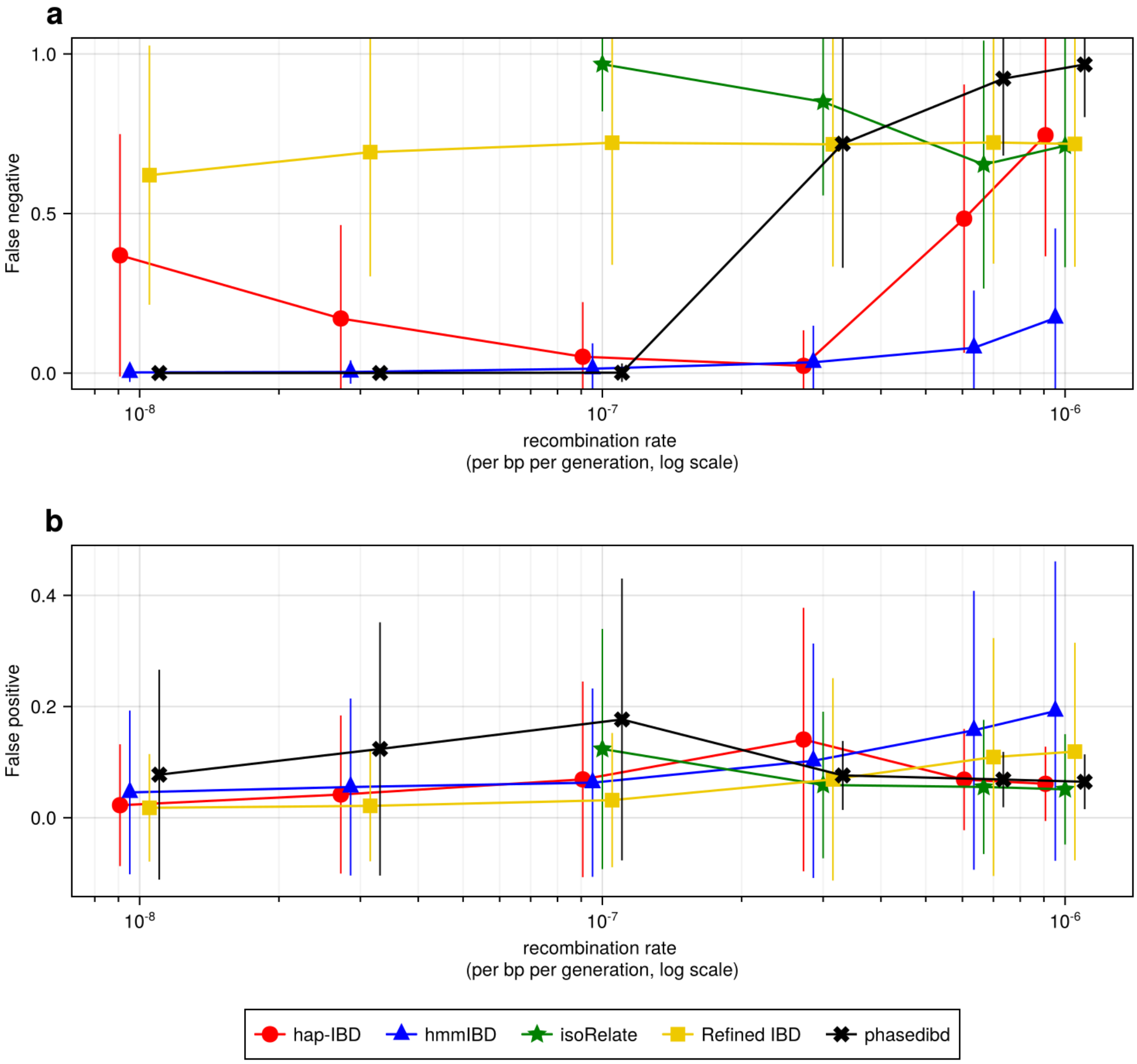

#### S2 Fig

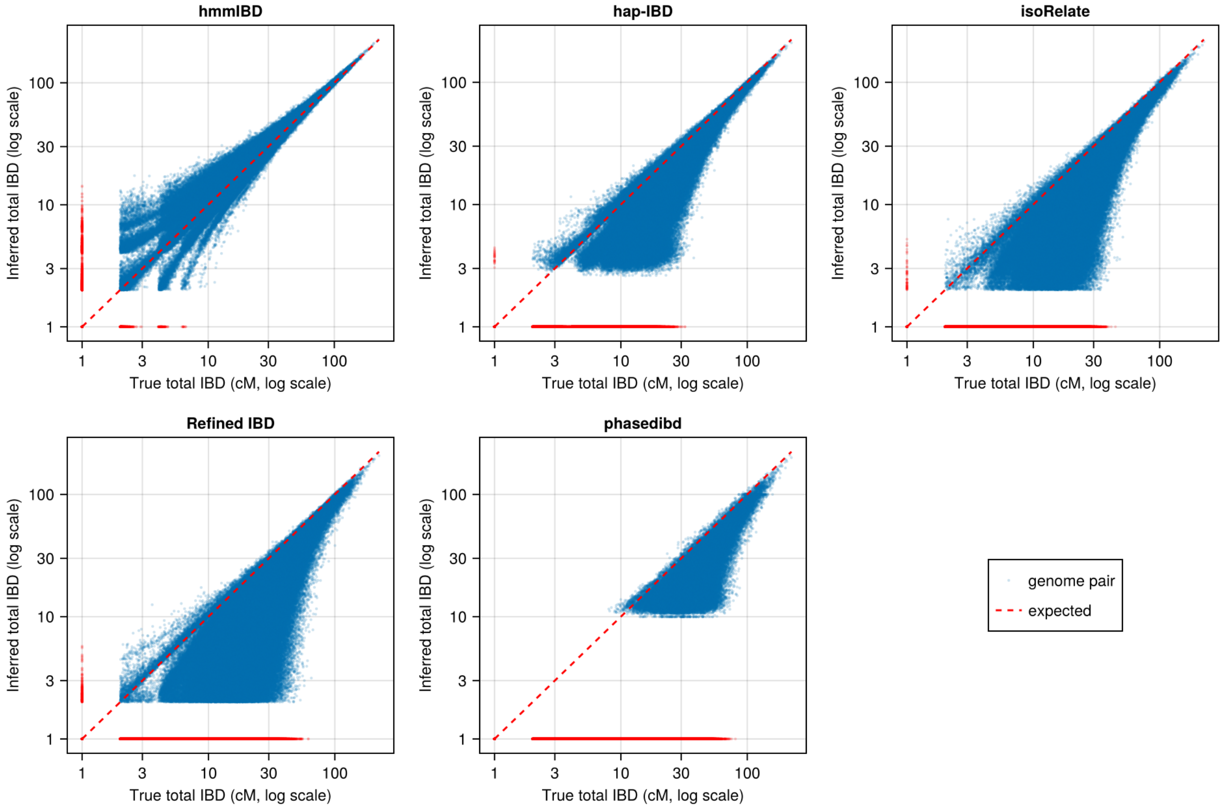

hmmIBD
hap-IBD
isoRelate
Refined IBD
phasedibd
Inferred total IBD (log scale)
True total IBD (cM, log scale)
1
3
10
30
100
genome pair
expected

#### S3 Fig

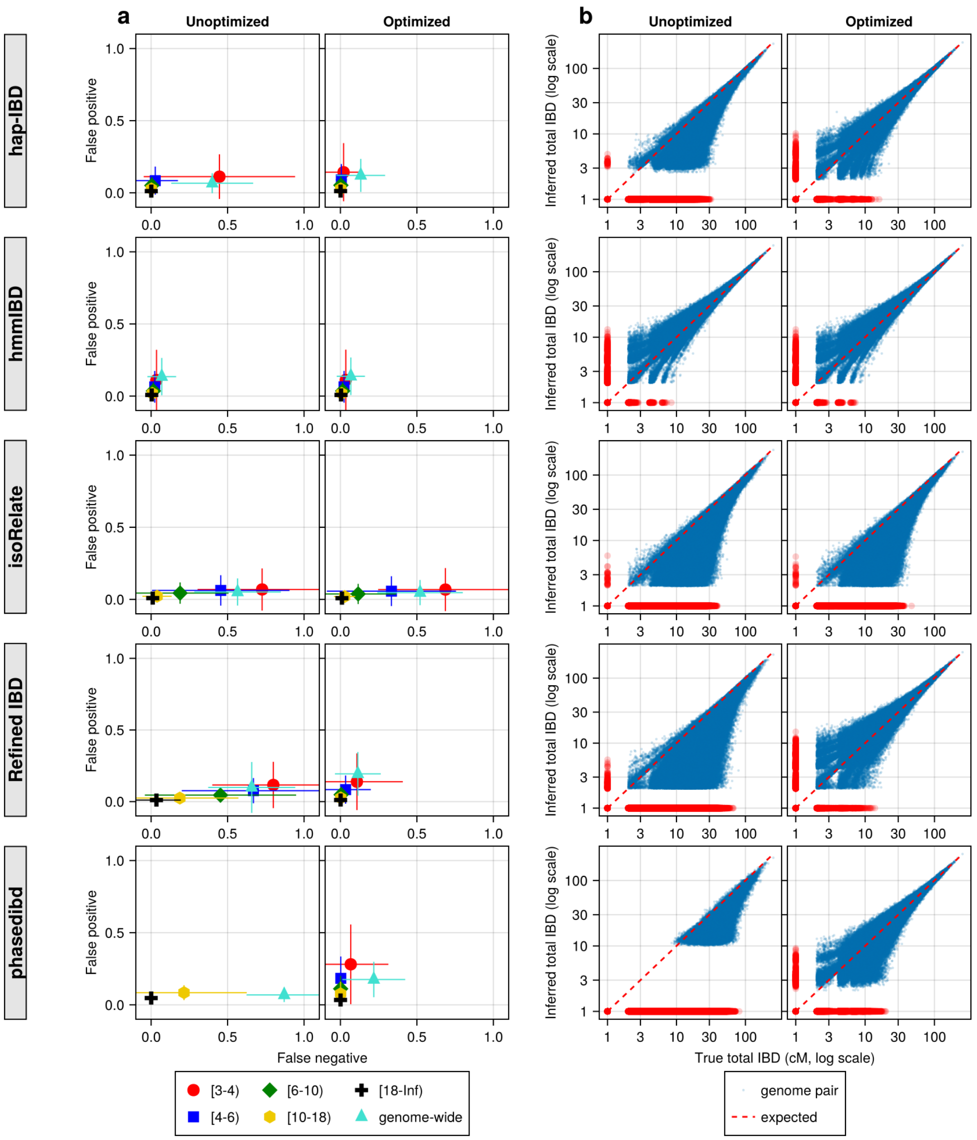

a
b
Unoptimized
Optimized
hap-IBD
hmmIBD
isoRelate
Refined IBD
phasedibd
False positive
False negative
Inferred total IBD (log scale)
True total IBD (cM, log scale)
[3-4)
[4-6)
[6-10)
[10-18)
[18-Inf)
genome-wide
genome pair
expected

#### S4 Fig

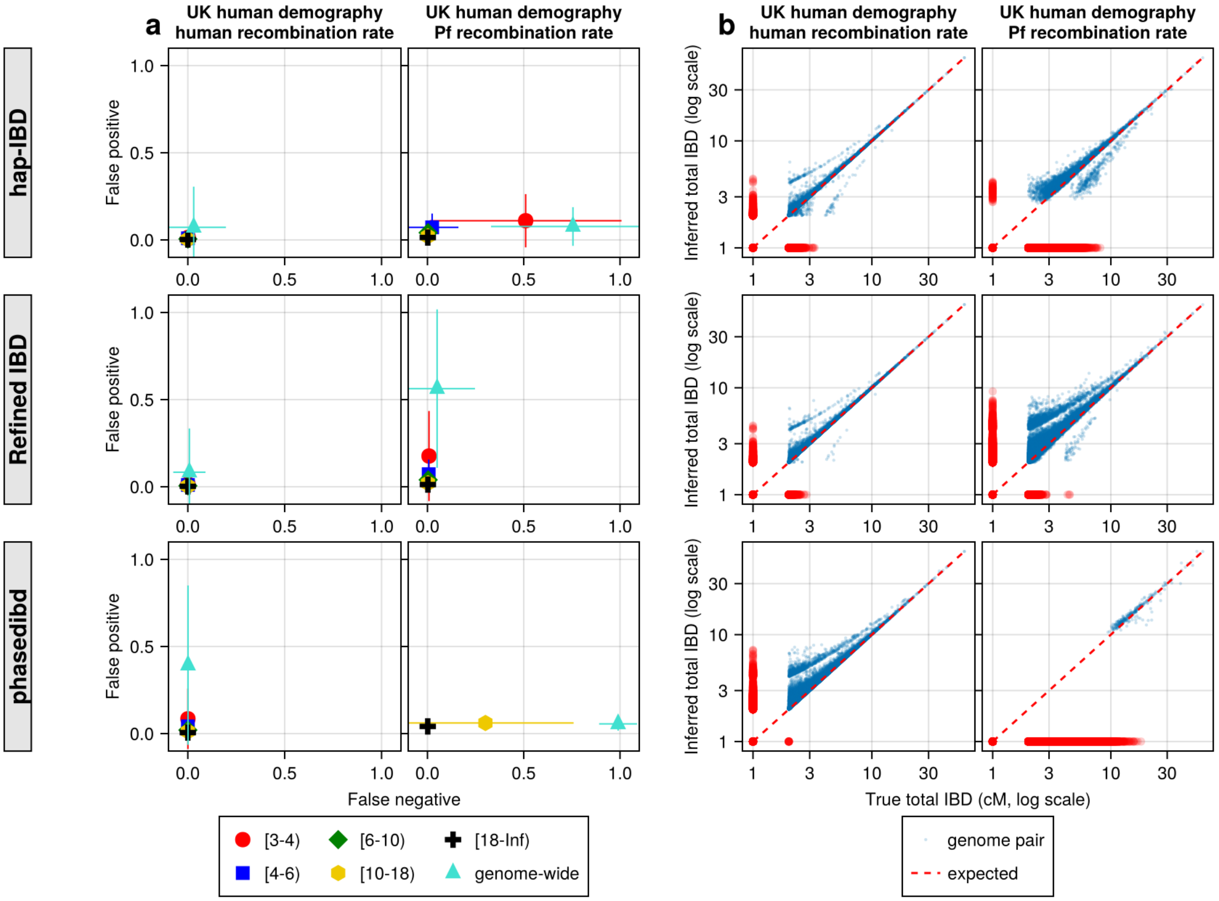

a
b
UK human demography human recombination rate
UK human demography Pf recombination rate
hap-IBD
Refined IBD
phasedibd
False positive
False negative
Inferred total IBD (log scale)
True total IBD (cM, log scale)
[3-4)
[4-6)
[6-10)
[10-18)
[18-Inf)
genome-wide
genome pair
expected


#### S5 Fig

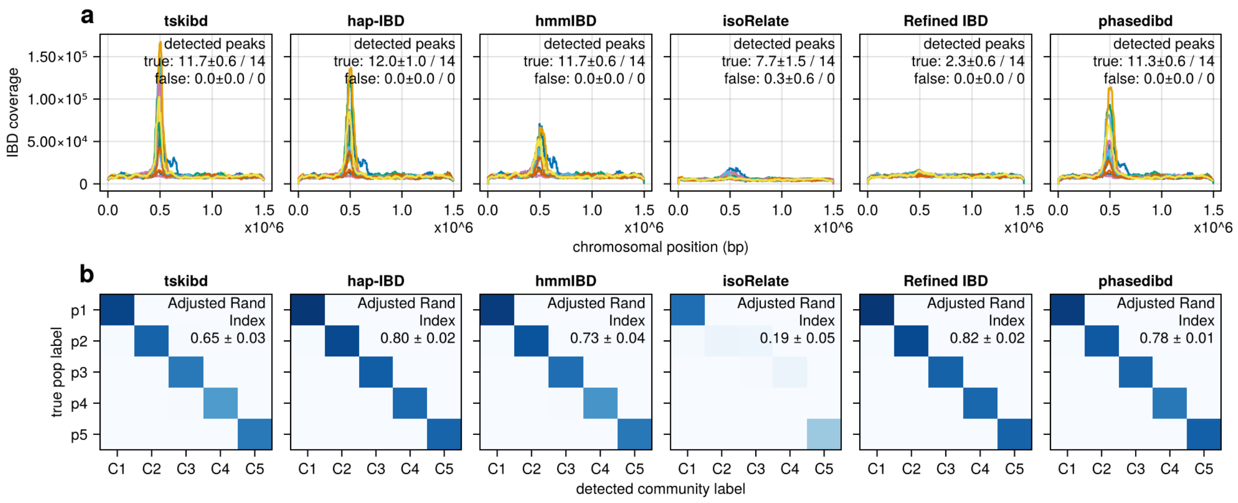

a
tskibd
hap-IBD
hmmIBD
isoRelate
Refined IBD
phasedibd
detected peaks
true: 11.7±0.6 / 14
false: 0.0±0.0 / 0
true: 12.0±1.0 / 14
false: 0.0±0.0 / 0
true: 11.7±0.6 / 14
false: 0.0±0.0 / 0
true: 7.7±1.5 / 14
false: 0.3±0.6 / 0
true: 2.3±0.6 / 14
false: 0.0±0.0 / 0
true: 11.3±0.6 / 14
false: 0.0±0.0 / 0
IBD coverage
chromosomal position (bp)
x10^6
b
Adjusted Rand Index
0.65 ± 0.03
0.80 ± 0.02
0.73 ± 0.04
0.19 ± 0.05
0.82 ± 0.02
0.78 ± 0.01
true pop label
p1
p2
p3
p4
p5
C1
C2
C3
C4
C5
detected community label

#### S6 Fig

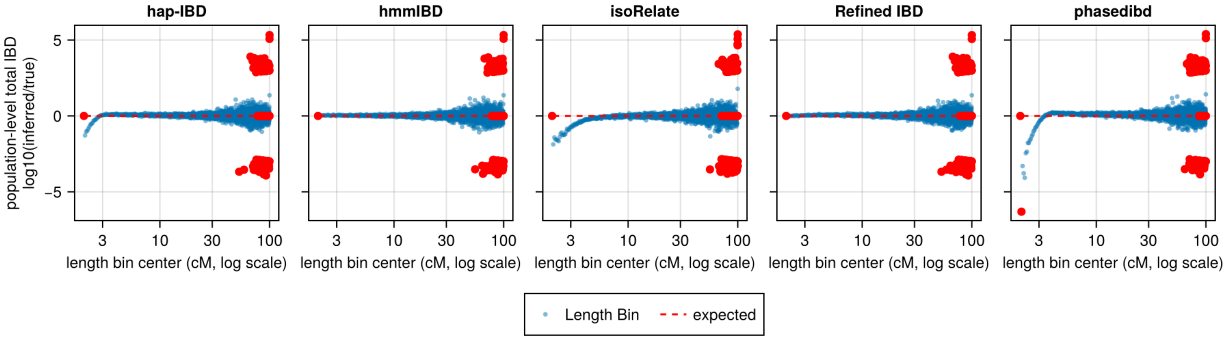

hap-IBD
hmmIBD
isoRelate
Refined IBD
phasedibd
population-level total IBD
log10(inferred/true)
5
0
−5
3
10
30
100
length bin center (cM, log scale)
Length Bin
expected


#### S7 Fig

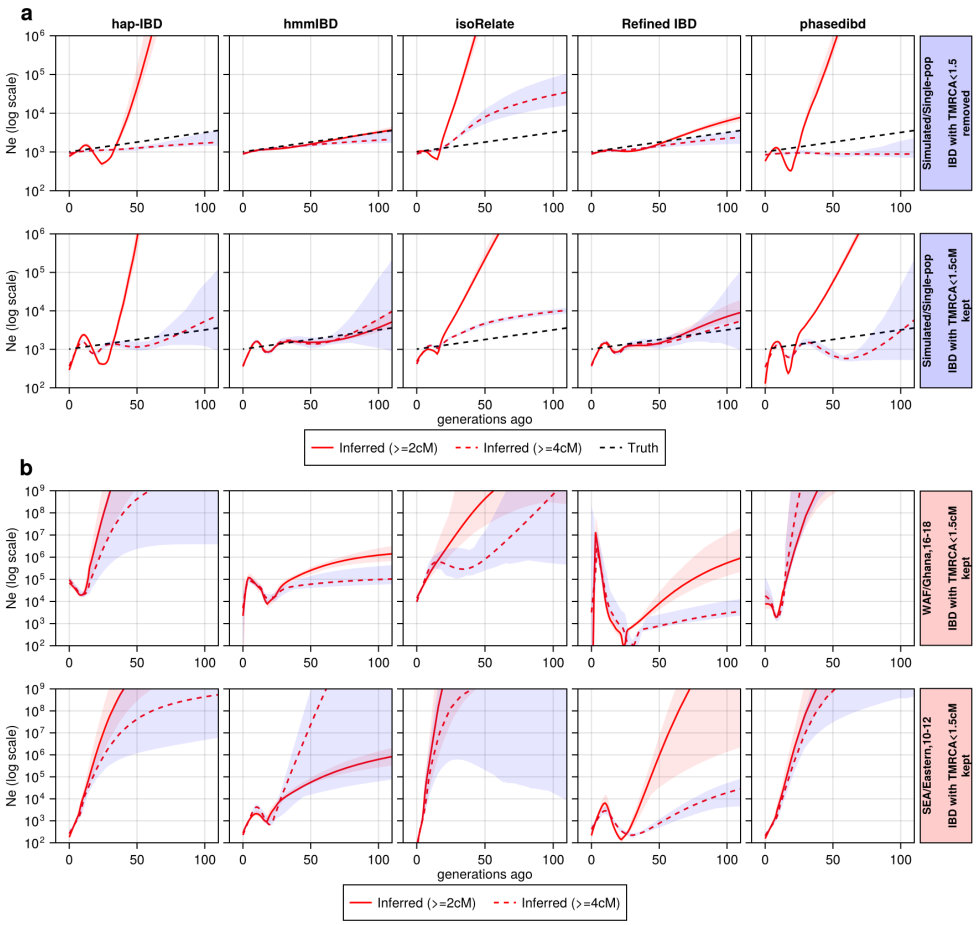

a
hap-IBD
hmmIBD
isoRelate
Refined IBD
phasedibd
Ne (log scale)
Simulated/Single-pop
IBD with TMRCA<1.5 removed
Simulated/Single-pop
IBD with TMRCA<1.5cM kept
generations ago
Inferred (>=2cM)
Inferred (>=4cM)
Truth
b
WAF/Ghana,16-18
IBD with TMRCA<1.5cM kept
SEA/Eastern,10-12
IBD with TMRCA<1.5cM kept
generations ago
Inferred (>=2cM)
Inferred (>=4cM)

#### S8 Fig

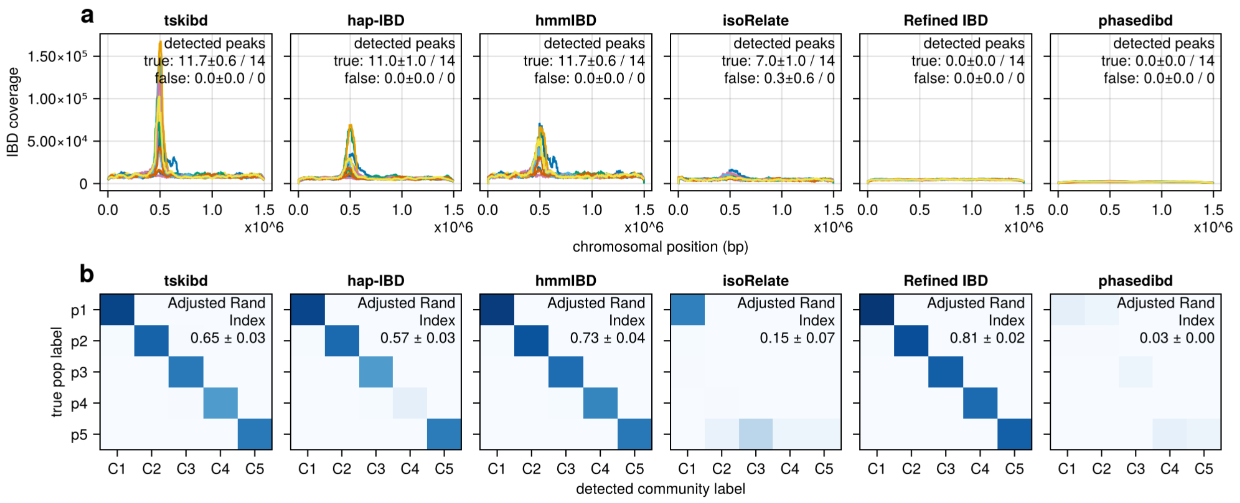

a
tskibd
hap-IBD
hmmIBD
isoRelate
Refined IBD
phasedibd
detected peaks
true: 11.7±0.6 / 14
false: 0.0±0.0 / 0
detected peaks
true: 11.0±1.0 / 14
false: 0.0±0.0 / 0
detected peaks
true: 11.7±0.6 / 14
false: 0.0±0.0 / 0
detected peaks
true: 7.0±1.0 / 14
false: 0.3±0.6 / 0
detected peaks
true: 0.0±0.0 / 14
false: 0.0±0.0 / 0
detected peaks
true: 0.0±0.0 / 14
false: 0.0±0.0 / 0
IBD coverage
1.50×10⁵
1.00×10⁵
5.00×10⁴
0
0.0
0.5
1.0
1.5
x10^6
chromosomal position (bp)
b
tskibd
hap-IBD
hmmIBD
isoRelate
Refined IBD
phasedibd
Adjusted Rand Index 0.65 ± 0.03
Adjusted Rand Index 0.57 ± 0.03
Adjusted Rand Index 0.73 ± 0.04
Adjusted Rand Index 0.15 ± 0.07
Adjusted Rand Index 0.81 ± 0.02
Adjusted Rand Index 0.03 ± 0.00
true pop label
p1
p2
p3
p4
p5
C1
C2
C3
C4
C5
detected community label


#### S9 Fig

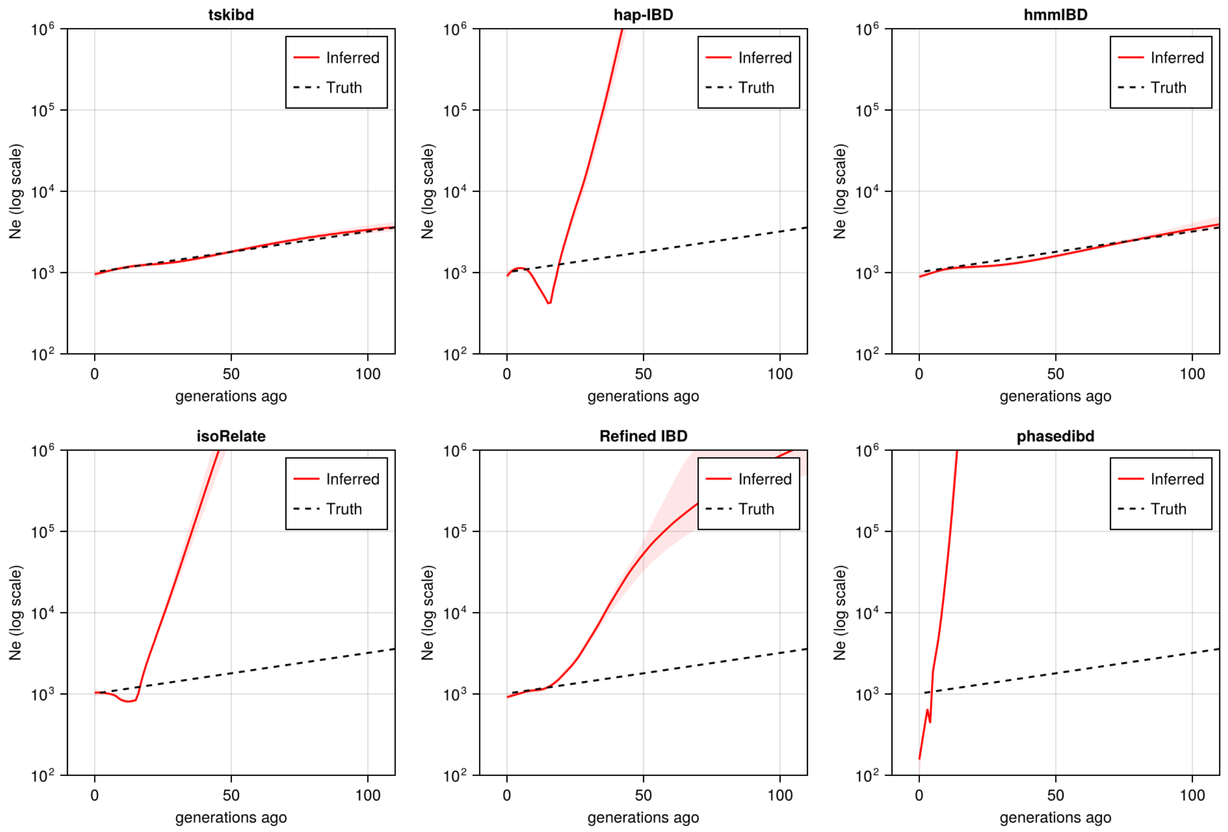

tskibd
hap-IBD
hmmIBD
isoRelate
Refined IBD
phasedibd
Inferred
Truth
Ne (log scale)
10⁶
10⁵
10⁴
10³
10²
0
50
100
generations ago

#### S10 Fig

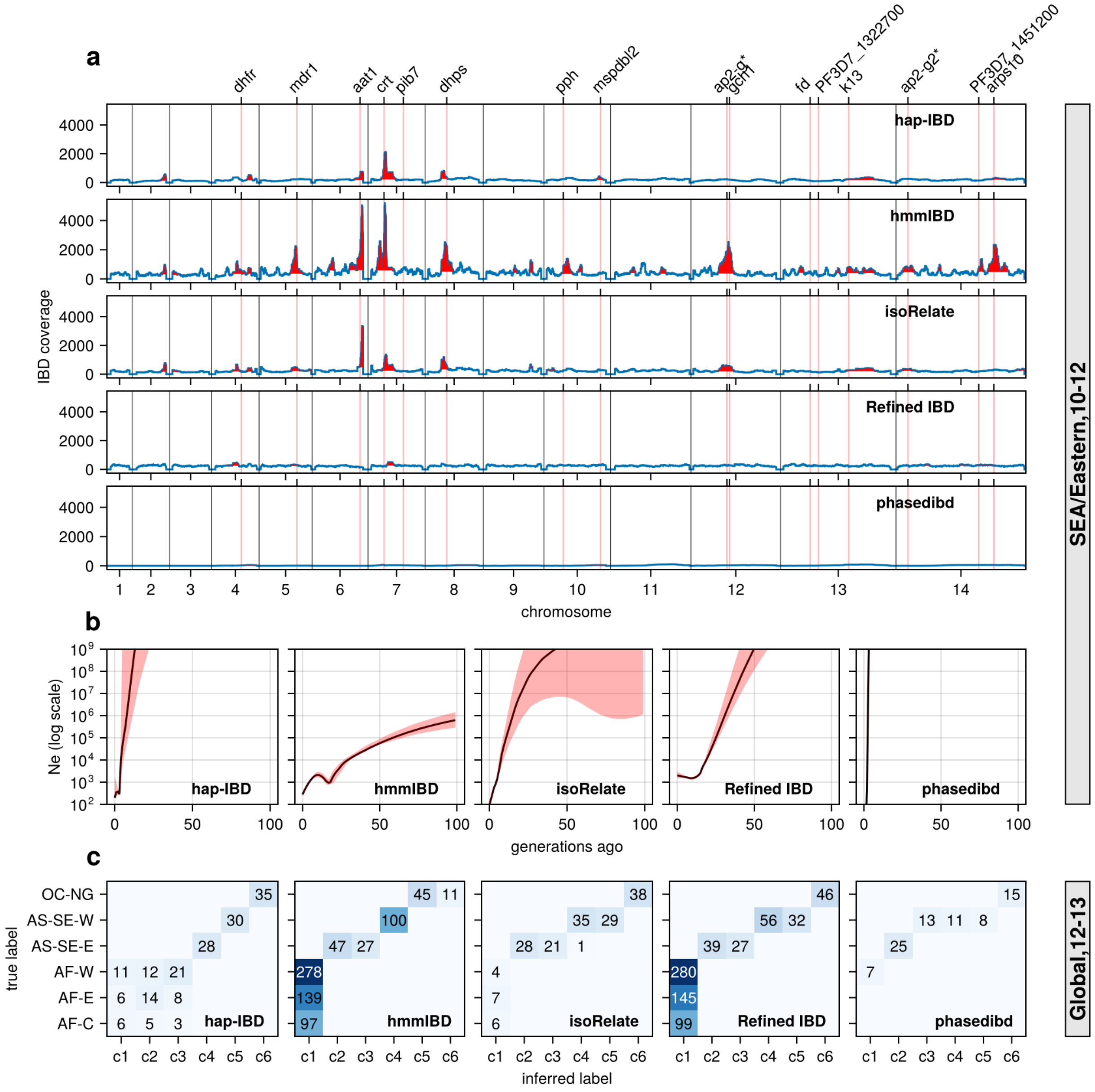

a
dhfr
mdr1
aat1
crt
pib7
dhps
pph
mspdbl2
ap2-g*
gch1
fd
PF3D7_1322700
k13
ap2-g2*
PF3D7_1451200
arps10
hap-IBD
hmmIBD
isoRelate
Refined IBD
phasedibd
IBD coverage
chromosome
SEA/Eastern,10-12
b
Ne (log scale)
generations ago
c
OC-NG
AS-SE-W
AS-SE-E
AF-W
AF-E
AF-C
true label
inferred label
c1 c2 c3 c4 c5 c6
Global,12-13

#### S11 Fig

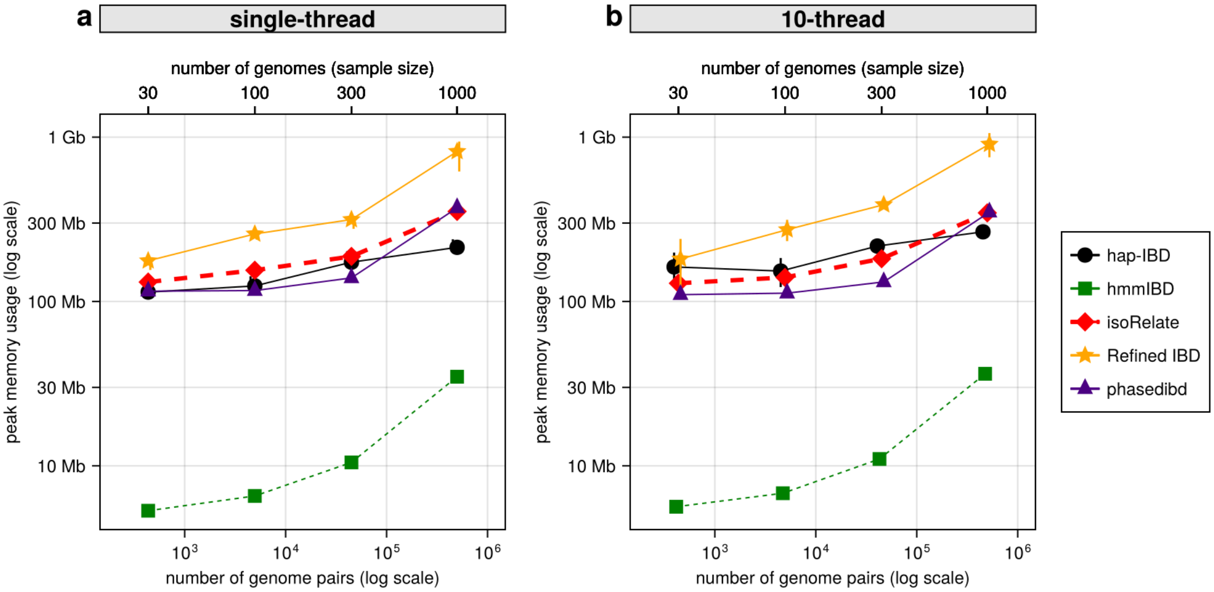

a
single-thread
b
10-thread
number of genomes (sample size)
30
100
300
1000
1 Gb
300 Mb
100 Mb
30 Mb
10 Mb
peak memory usage (log scale)
$10^3$
$10^4$
$10^5$
$10^6$
number of genome pairs (log scale)
hap-IBD
hmmIBD
isoRelate
Refined IBD
phasedibd

### 2 Supplementary Tables

#### S1 Table

| IBD caller | Program parameter | Default value | Optimized/used value |
|---|---|---|---|
| `hap-IBD`<br>• version: 1.0 23Apr20.f1a<br>• Browning *et al.* 2020 | min-output | 2.0 | mincm * |
| | min-seed | 2.0 | mincm |
| | min-extend | 1.0 | 1.0 |
| | max-gap | 1000 | 1000 |
| | min-markers | 100 | 70 |
| `hmmIBD`<br>• github.com/glipsnort/hmmIBD<br>• Commit: a2f796e<br>• Schaffner *et al.* 2018 | rec_rate (in hmmIBD.c) | $7.4 \times 10^{-7}$ | $6.67 \times 10^{-7}$ |
| | m | 5 | 5 |
| | n | no limit | 100 |
| `isoRelate`<br>• github.com/bahlolab/isoRelate<br>• Commit: 109ee47<br>• Henden *et al.* 2018 | isolate.max.missing | 0.1 | imputed |
| | snp.max.missing | 0.1 | imputed |
| | maf | 0.01 | 0.1 |
| | minimum.length.bp | 50,000 | mincm $\times$ 15,000 |
| | minsnp | 20 | 20 |
| `Refined IBD`<br>• version: 17Jan20.102<br>• Browning *et al.* 2013 | length | 1.5 | mincm |
| | lod | 3.0 | 1.6 |
| | scale | data-dep | data-dep |
| | window | 40 | 40 |
| | trim | 0.15 | 0.15 |
| `phased IBD`<br>• github.com/23andMe/phasedibd<br>• Commit: 9a7b949<br>• Freyman *et al.* 2021 | template | $\begin{bmatrix} 1 & 0 & 1 & 0 \\ 0 & 1 & 0 & 1 \\ 1 & 1 & 0 & 0 \\ 0 & 0 & 1 & 1 \\ 1 & 0 & 0 & 1 \\ 0 & 1 & 1 & 0 \end{bmatrix}$ | $\begin{bmatrix} 0 & 1 & 1 & 1 \\ 1 & 0 & 1 & 1 \\ 1 & 1 & 0 & 1 \\ 1 & 1 & 1 & 0 \end{bmatrix}$ |
| | L_m | 300 | 80 |
| | L_f | 3.0 | mincm * |

#### S2 Table

| IBD caller | Program parameter | Tested values | Comment |
|---|---|---|---|
| `hap-IBD` | max-gap | (3, 30, 100, 300, 1000) | little to no effect |
| | min-marker | coarse: (3, 10, 30, 100)<br>finetune:(30, 40, 50, 60, 70, 80, 100) | optimal around 70 |
| `hmmIBD` | m | (2, 5, 10) | little to no effect |
| | n | (10, 30, 100, 300, Inf) | little to no effect |
| | min-maf | (0.001, 0.01) | little to no effect |
| `isoRelate` | min-maf | (0.01, 0.03, 0.1) | 0.1 reduces FN for longer IBD |
| | min-snp | (1, 3, 10, 15, 20, 40, 80, 160) | $\leq$ 40 reduces FN |
| `Refined IBD` | min-maf | (0.01, 0.1) | little to no effect |
| | lod | (1.1, 1.2, 1.4, 1.6, 1.8, 2, 3, 4, 8) | lowest FN at 1.6 |
| | window | (20, 40) | little to no effect |
| | trim | (0.01, 0.02, 0.05, 0.08, 0.10, 0.12, 0.15) | little to no effect |
| `phased IBD` | template | tolerate 1 or 2 mismatch in every 4 SNPs | optimal to tolerate 1 mismatch |
| | L__m | (50, 80, 90, 100, 110, 130, 150, 200, 250, 300) | optimal around 80 |
| | min-maf | (0.001, 0.01, 0.1) | optimal around 0.01 |

#### S3 Table

| Population | 2012 | 2013 | Total |
|---|---|---|---|
| AF-C | 66 | 37 | 103 |
| AF-E | 58 | 139 | 197 |
| AF-W | 27 | 273 | 300 |
| AS-SE-E | 75 | 25 | 100 |
| AS-SE-W | 85 | 59 | 144 |
| OC-NG | 17 | 40 | 57 |
| Total | 328 | 573 | 901 |

#### S4 Table

| Country | 2010 | 2011 | 2012 | Total |
|---|---|---|---|---|
| Cambodia | 61 | 82 | 43 | 186 |
| Laos | 27 | 27 | 15 | 69 |
| Thailand | 0 | 0 | 1 | 1 |
| Vietnam | 28 | 44 | 16 | 88 |
| Total | 116 | 153 | 75 | 344 |

#### S5 Table

| Admin level 1 | 2016 | 2017 | 2018 | Total |
|---|---|---|---|---|
| Eastern | 0 | 0 | 6 | 6 |
| Greater Accra | 14 | 13 | 57 | 84 |
| Upper East | 63 | 199 | 212 | 474 |
| Volta | 21 | 0 | 0 | 21 |
| Total | 98 | 212 | 275 | 585 |

### 3 Supplementary Data

#### S2 Data

a, `hap-IBD`, single population model, *Pf* recombination rate.

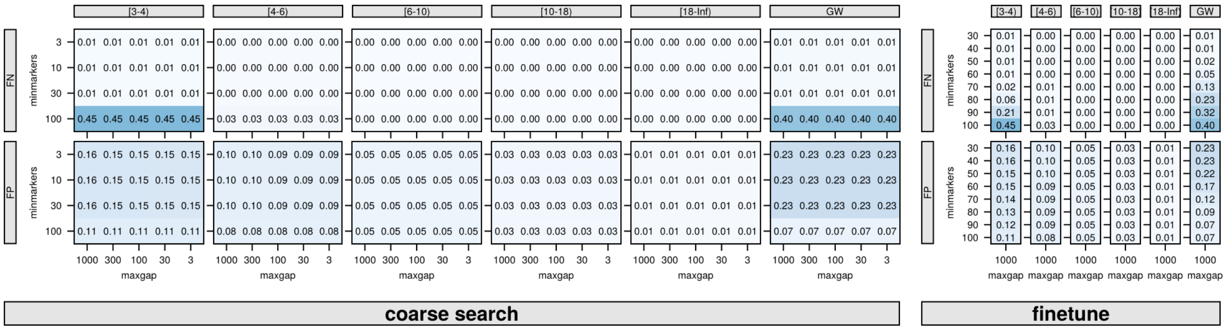


b, `hap-IBD`, multiple population model, *Pf* recombination rate.

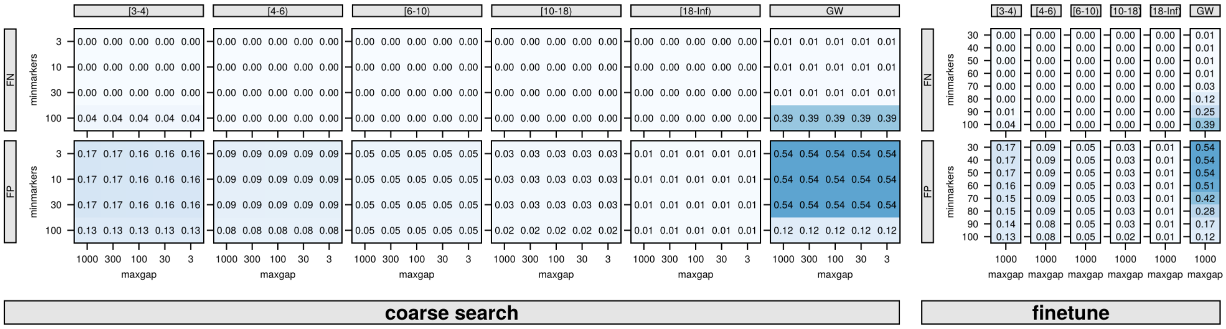


c, `hap-IBD`, UK human population model, Human recombination rate.

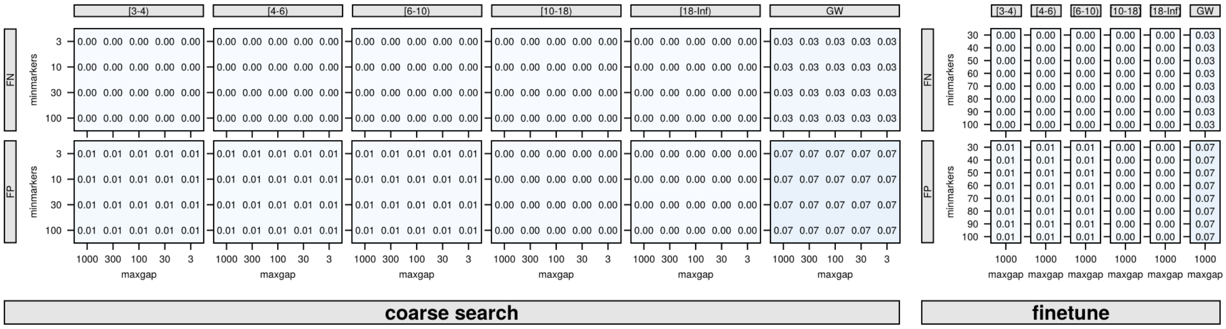

d, `hap-IBD`, UK human population model, *Pf* recombination rate.

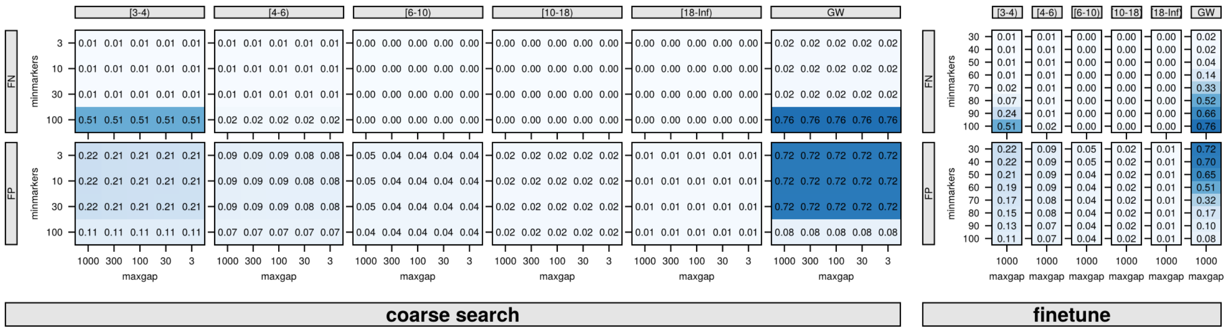


e, `hmmIBD`, single population model, *Pf* recombination rate.

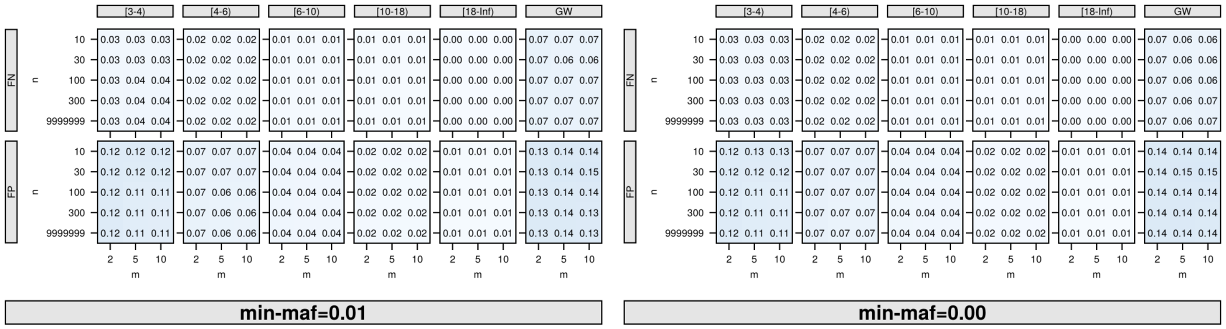


f, `hmmIBD`, multiple population model, *Pf* recombination rate.

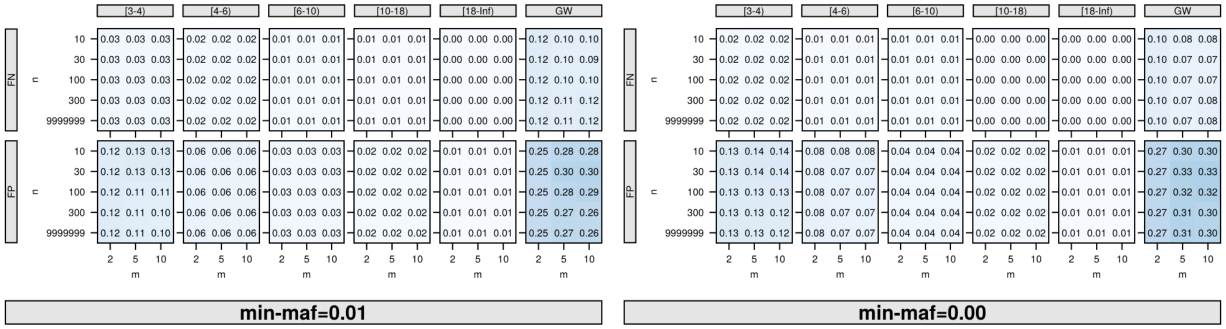


g, `hmmIBD`, UK human population model, *Pf* recombination rate.

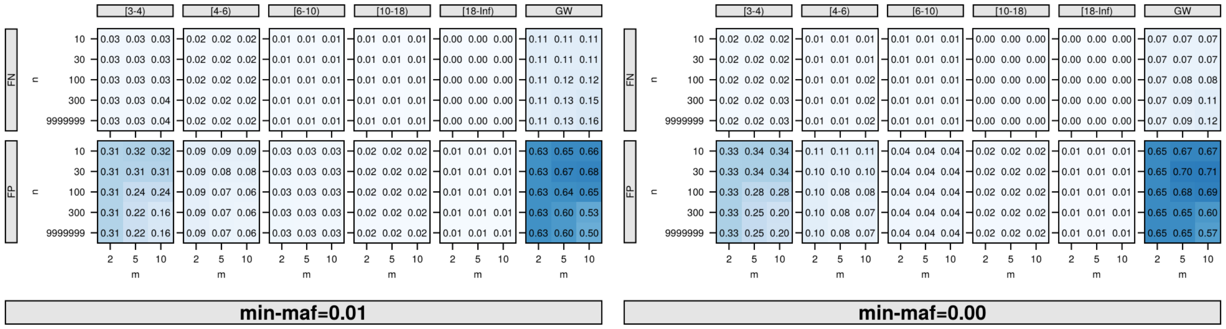

h, `isoRelate`, single population model, *Pf* recombination rate.

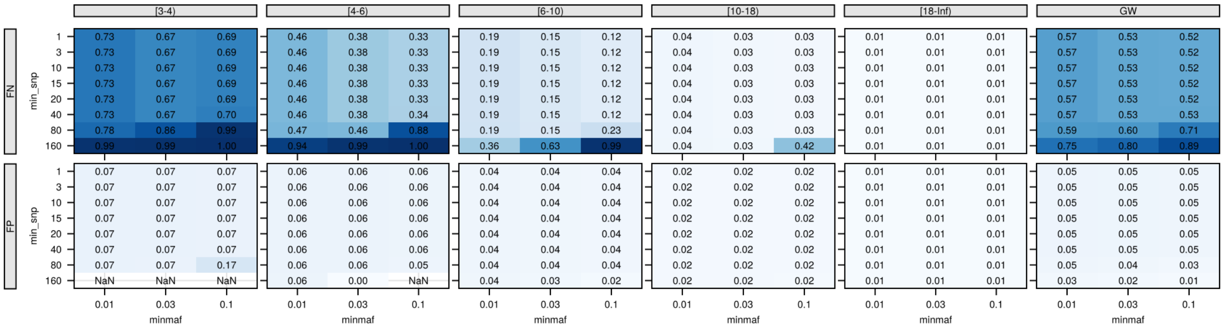


i, `isoRelate`, multiple population model, *Pf* recombination rate.

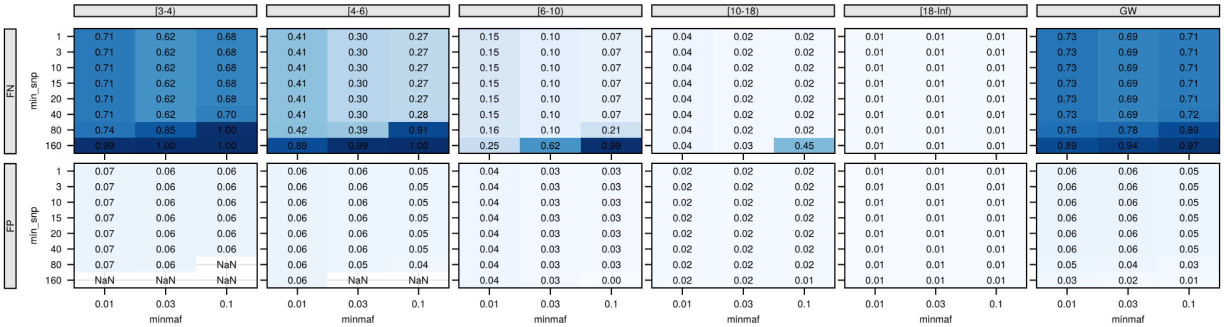


j, `isoRelate`, UK human population model, *Pf* recombination rate.

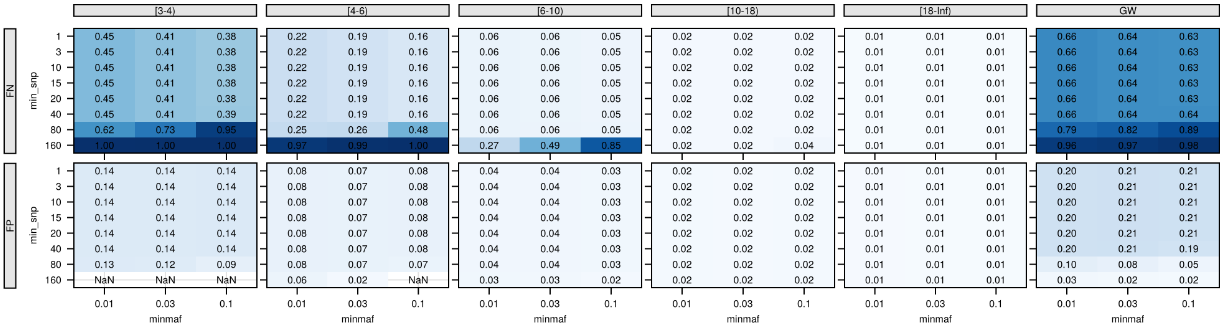


k, `Refined IBD`, single population model, *Pf* recombination rate.

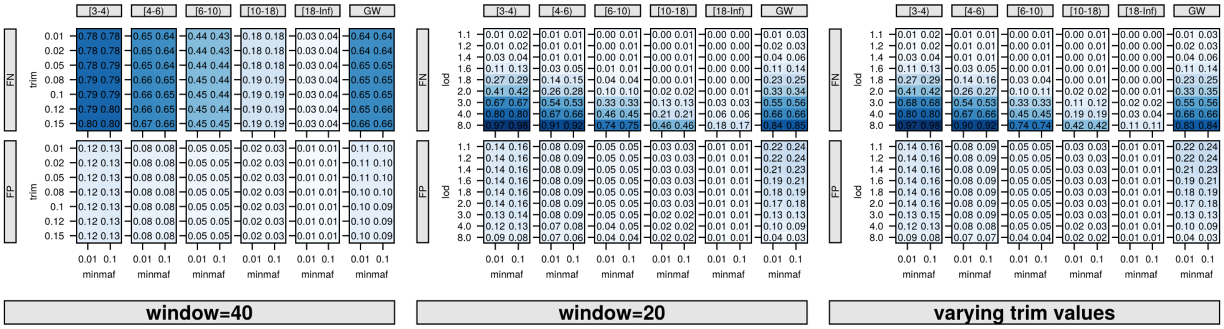

l, `Refined IBD`, multiple population model, *Pf* recombination rate.

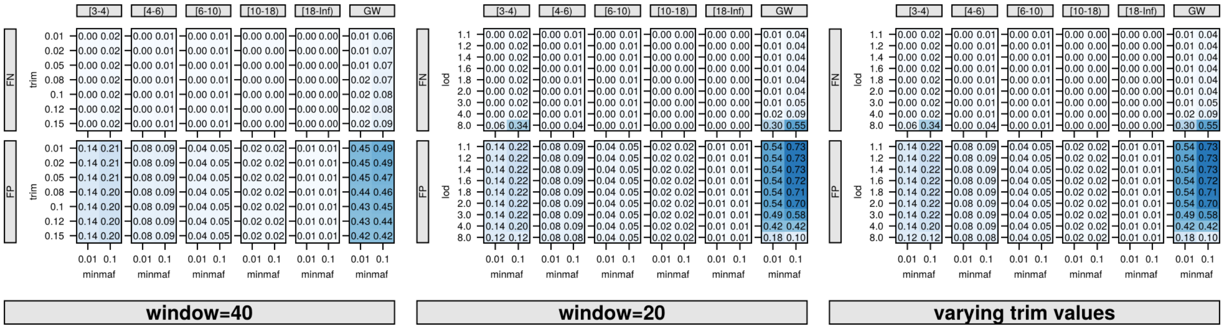


m, `Refined IBD`, UK human population model, Human recombination rate.

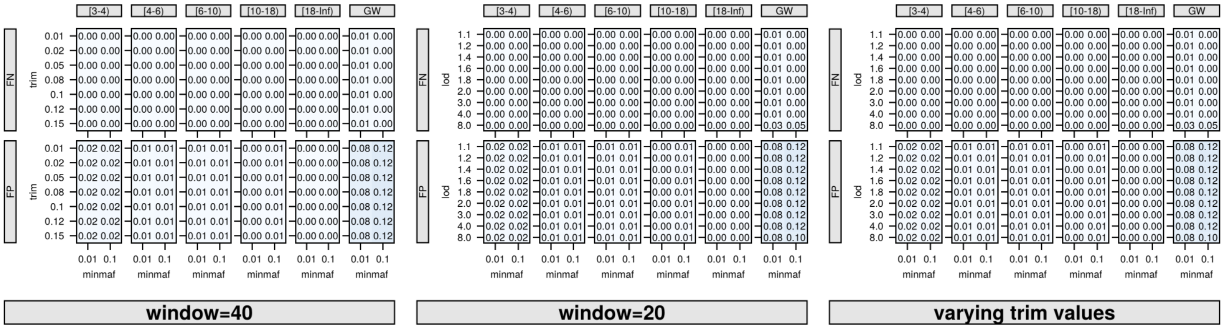


n, `Refined IBD`, UK human population model, *Pf* recombination rate.

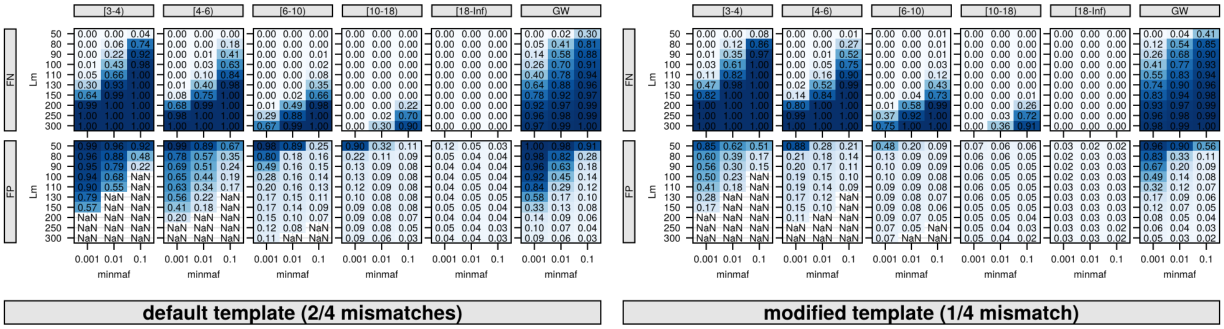


o, `phased IBD`, single population model, *Pf* recombination rate.

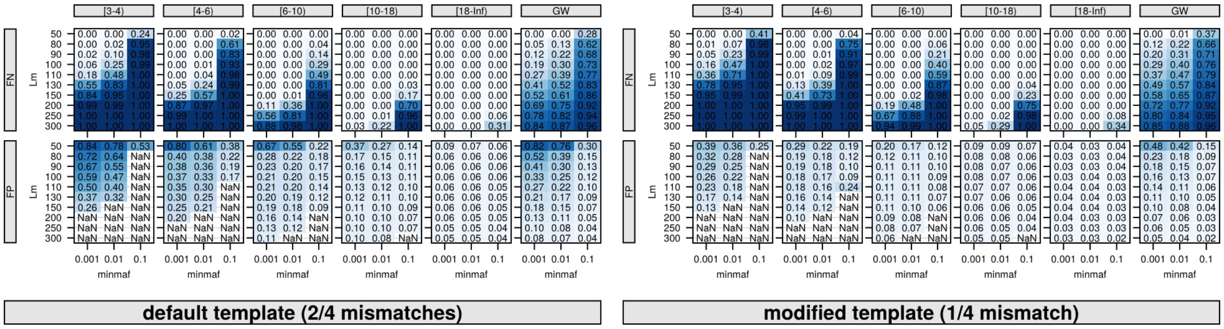

p, `phased IBD`, multiple population model, *Pf* recombination rate.

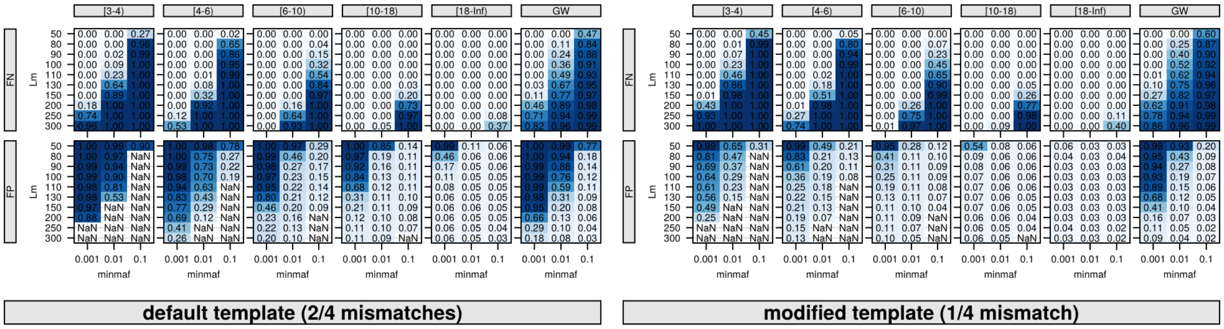


q, `phased IBD`, UK human population model, Human recombination rate.

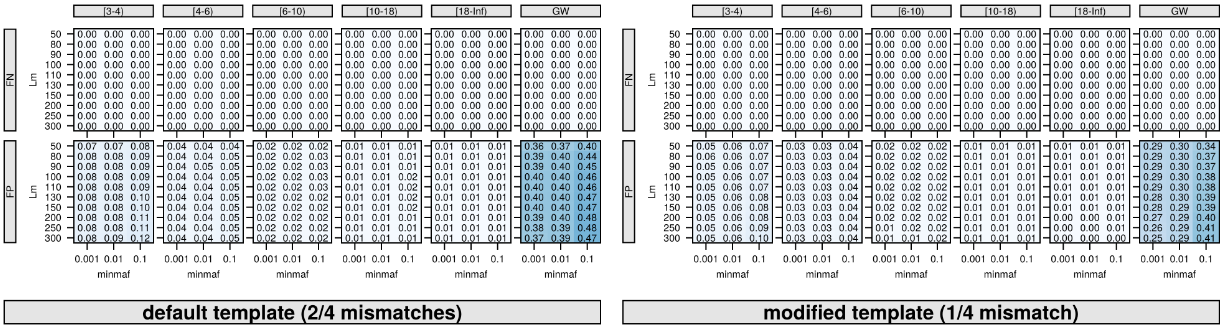


r, `phased IBD`, UK human population model, *Pf* recombination rate.

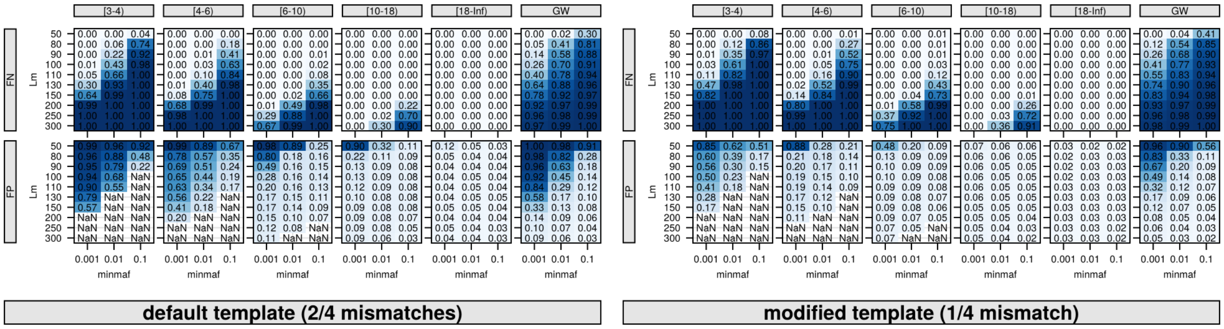
